# “Investigating the effect of obesity on adipose-derived stem cells (ASCs) using Göttingen Minipigs”

**DOI:** 10.1101/2022.02.11.477943

**Authors:** Maria Meyhoff-Madsen, Esben Østrup, Merete Fredholm, Susanna Cirera

## Abstract

Obesity is associated with low grade inflammation, which may adversely impact the biological functions of adipose tissue and consequently of adipose-derived stem cells (ASCs). Studies in humans and rodents have described that obesity alters ASC properties and functionality, compromising their therapeutic prospects. The Göttingen Minipig (GM) is a commonly used obesity model. Nevertheless, there are no studies investigating the effect of obesity on ASCs from GM, which could constitute a valuable addition to both obesity modelling and adult stem cells investigations.

In this study, we isolated subcutaneous ASCs from lean and obese GM to investigate the effect of obesity on cell behavior and differentiation capacity. During culturing, we observed an inherent difference in cell morphology between lean and obese ASCs. Upon adipogenic induction, obese-ASCs readily differentiated, developing significantly larger amounts of adipocytes than corresponding lean-ASCs, hinting at a predisposition towards adipogenic differentiation. Expression profiling of obesity-related genes in cell cultures, before and after adipogenic differentiation, revealed a tendency towards up-regulation in differentiated obese-cultures. Altogether, our results indicate that stem cells from obese donors could display different therapeutic properties. In summary, our results point towards GM as a valuable model for future ASCs investigations in healthy and obese states.

## Introduction

Mesenchymal stem/stromal cells (MSCs) are self-renewing, multipotent cells, distinguished from other stem cell types by plastic-adherence, adipocyte, osteoblast and chondroblast differentiation capacity, and expression of specific surface proteins [1]. MSCs represent an increasing prospect in e.g. regenerative medicine, due to their ability to differentiate to a multitude of tissues and conform to tissue phenotype and activity *in vivo* [2–5]. MSCs are usually extracted from bone marrow, which is a painful procedure and results in low cell numbers. A promising alternative is the adipose tissue which constitutes a useful source of MSCs (Adipose derived stem cells or ASCs) as it allows easy harvesting, isolation and culturing, larger yields, less donor-site morbidity and more rapid proliferation than bone marrow-derived MSCs [5, 6].

Obesity is a growing problem worldwide, constituting a major risk factor for developing comorbidities such as diabetes, cardiovascular diseases and cancer [7]. Obesity is defined as an enlargement of adipose tissue by hyperplasia (increase in cell number) and/or by hypertrophy (increase in cell size) [8]. Adipocyte hyperplasia is due to activation of multipotent stem cells (ASCs) that will generate new adipocytes to sustain the elevated demand for fat storage. Indeed, subcutaneous adipose tissue (SAT) is the most important fat depot characterized by its capacity to expand in response to surplus of energy. Altered adipogenesis in SAT leads to dysfunctional, hypertrophic adipocytes together with metabolic complications associated to obesity (i.e chronic low-grade inflammation and insulin resistance) [9]. In fact, an increasing number of reports have described obesity to cause alterations in ASCs properties [10–17], which might affect their therapeutic prospects in regenerative cell-based therapies. On the other hand it has also been proposed that the preservation of the functionality of subcutaneous adipocyte precursors could contribute to some obese individuals remaining metabolically healthy (MHO) [9].

However, since the use of autologous ASCs is associated with a lower probability of rejection or extrusion [18] it is important to consider individual donor characteristics (i.e. obesity and metabolic status) when using ASCs for therapeutic applications.

Relevant animal disease models are of fundamental importance to study the properties and potential of stem cells for possible future applications in human medicine. In this context, the pig offers a good translational model for human disease studies due to similar physiology, body fat distribution, metabolism, dietary habits, immunology and genome to humans [19, 20]. Zhu et al. [18] investigated if obesity would affect ASC function using domestic pigs and found that *in vitro* ASCs from obese pigs showed enhanced adipogenic and osteogenic differentiation. Göttingen Minipig (GM) is a small-size pig breed prone to obesity when fed *ad libitum*. Interventions with diets containing excess fat and sugar in GM have resulted in symptoms similar to human obesity; i.e. insulin resistance, impaired glucose tolerance, increased plasma cholesterol, triglyceride, free fatty acid levels and adipose tissue inflammation in pigs [21–25]. Moreover, diabetes-like symptoms can be induced in pigs [22, 25], making the GM an often used model in studies of obesity, metabolic syndrome and diabetes. Additionally, some studies have indicated that the observed metabolic diseases in GM could be dietary induced, and that GM, when fed standard pig chow with no excess fat and sugar, display a healthy lipid profile and insulin sensitivity, despite being obese mimicking the metabolic healthy obese (MHO) phenotype in humans [22, 26–28].

To our knowledge, no investigations have been made on the effect of obesity on ASCs from GM, which would however constitute a valuable addition to both obesity modelling and stem cell knowledge. The aim of this study was to investigate the possible effect of obesity status of the donor on cell behavior and differentiation capacity of ASC derived from lean and obese subcutaneous adipose tissue (SAT) from GM.

## Materials and methods

### Tissue isolation

Lean subcutaneous adipose tissue (SAT) samples were sampled from the neck region of castrated male Göttingen Minipigs at 9 months of age (N=3) after euthanasia, and cooled on ice during transportation to our lab facilities. Pigs were fed a standard SDS minipig diet and weighting 16.7-18.6 kg at time of euthanasia.

Obese SAT samples were sampled from the same regions as above from 9.5 months old ovariectomized obese female Göttingen Minipigs (N=3) during surgical anesthesia (zoletil mixture, IM) and cooled on ice during transportation to our lab facilities. Pigs were fed a high fat diet (Altromin 9033) and weighed 50.6-54 kg at time of sampling and were qualified as obese.

### Cell isolation

Upon arrival to the lab facilities, SAT was cut in 2-4mm pieces, mixed with 0.2% collagenase IV (Sigma-Aldrich) and incubated in an incubator shaker at 37°C, 50-80rpm for >3hours. Liquid was transferred to a Falcon tube and centrifuged at 300g for 10min. Supernatant was removed and pellet was resuspended in PBS (Sigma-Aldrich) and filtered through a 70μm filter retainer (Sigma-Aldrich). Subsequently, samples were centrifuged at 300g for 10min. and supernatant discarded. Pellet was solubilized in 1ml Dulbecco’s Modified Eagle’s Medium/Nutrient Mixture F-12 Ham (DMEM/F-12)(Sigma-Aldrich) with 20% Fetal bovine serum (FBS)(Gibco) and 5% Dimethyl Sulfoxide (DMSO)(Sigma-Aldrich) and frozen at - 80°C for >24hours and subsequently moved to liquid nitrogen for long-term storage.

### Cell culturing

Samples were thawed by addition of 38.5°C thawing media comprising DMEM/F-12 (Sigma-Aldrich) + 20% FBS (Gibco) + 1% Penicillin-Streptomycin (Thermo Fisher Scientific). For samples with DMSO content >0.5% a centrifugation step was added: 300g for 8min., upon which media was removed and pellet dissolved in new thawing media. Media was distributed to T25/T75 flasks and incubated at 38.5°C, 5% CO_2_. Media was changed every 2-3 days for new media: DMEM/F-12 (Sigma-Aldrich) + 10% FBS (Gibco) + 1% Penicillin-Streptomycin (Thermo Fisher Scientific). 1% Amphotericin B (Fungizone)(Sigma-Aldrich) was added to media until 1st passage. Cells were passaged when reaching >80% confluence or dense cell islands had formed in stromal vascular fraction (SVF) non-passaged (P0) cultures. In passaging, media was removed to a 50 ml Flacon tubes and cells were washed with PBS. Trypsin-EDTA (0.25%), phenol red (Sigma-Aldrich) was added and samples were incubated more than10min at 38.5°C until cells were floating in suspension. Trypsin was then neutralized by re-addition of collected media and the mixture was transferred to a 50 ml Falcon tube and centrifuged at 300g for 8min. Media was then removed and pellet resuspended in growth or freezing media. Cell numbers and viability were estimated using a Countess™ II FL Automated Cell Counter (Invitrogen) according to manufacturer’s instructions. We had previously tested culturing of 5000 cells/cm2 and 10,000cells/cm2 for 2 passages. Cells seeded at 10,000cells/cm2 had higher grow rate and showed less senescence and therefore, this condition was used in our experiments. After 2 passages, cells were frozen in DMEM/F-12 (Sigma-Aldrich) + 20% FBS (Gibco) + 5% DMSO (Sigma-Aldrich) for >24 hours in a freezing container before transfer to liquid nitrogen. Before freezing, contamination was discovered in obese sample F2 and it was excluded from further experiments.

### Differentiation

Specific media were used for *in vitro* differentiation of the ASCs into adipogenic, osteogenic and chondrogenic cells respectively. Prior to adipogenic differentiation, 2 different adipogenic media were tested, as described by Kim *et al*. (2016) and Zimmerlin *et al*. (2010) [29, 30] (see Supplementary text for a detailed description of each media).

Upon reaching confluence, cells were trypsinized and distributed to well plates in concentrations of 10,000cells/cm^2^. Media was changed for new growth media every 2-3 days. Upon reaching confluence, growth media was replaced with adipogenic, osteogenic or growth media (control). Cells were incubated at 38.5°C, 5% CO_2_, with media changed every 2-3 days for 21 days. For chondrogenic differentiation 250,000 cells were added to 600-650μl chondrogenic medium (total 700μl) in 15ml Falcon tubes and centrifuged at 600g for 5min. Lids of Falcon tubes were loosened to allow air flow, and samples were incubated at 38.5°C, 5% CO_2_. Chondrogenic medium was changed every 2-3 days, by removal of 0.5ml medium, without disturbing the pellet, and addition of 0.5ml new chondrogenic medium for 21 days.

At day 21, adipogenic samples were lysed in TRI Reagent (MRC) and the lysate was stored at −80°C until RNA extraction. For staining procedures, media was removed and ~0.5ml/cm^2^ 4% Formalin in PBS was added to each well or tube, and samples were stored at 4°C for 3-5 weeks. One week before staining, formalin was replaced by PBS. Cultures were stained with Oil Red O (Sigma-Aldrich) (for adipogenic differentiation), Alizarin Red S (Sigma-Aldrich) (for osteogenic differentiation), or Alcian Blue pH=2.5 (Sigma-Aldrich), cast in Tissue-Tek (Sakura® Finetek) and sliced in 20μm cuts in a CM1950 cryomicrotome (for chondrogenic differentiation).

After imaging, Oil Red O was eluded, as described in [31], and absorption measured in duplicate at 490nm in a clear 96-well polystyrene microtiter plate on a Powerwave X 340 Microplate Scanning Spectrophotometer (BIO-TEK).

### RNA isolation and cDNA synthesis

RNA from cells was purified following standardized TRI Reagent protocol. Samples were DNase-treated using column-based clean-up protocol (Qiagen). Concentration and integrity were estimated on a NanoDrop 1000 spectrophotometer and Bioanalyzer 2100 (Agilent Technologies) using a RNA 6000 Nano Kit (Agilent Technologies), respectively. cDNA was synthesized in duplicates from 100 ng total RNA using ImProm II reverse transcriptase (Promega), RNAsin (Promega) and 3:1 mixture of random hexamers/OligodT (Sigma-Aldrich), following manufacturer’s instructions. cDNA samples were 8 times diluted before used in quantitative real-time (qPCR) experiments.

### qPCR gene profiling

Four adipogenesis-related genes were profiled by qPCR in cell cultures before adipogenic induction (T0) and 21 days after differentiation (T21) in obese and lean samples. These genes are: Adiponectin *(ADIPOQ)*, Klotho Beta *(KLB)*, Lipoprotein Lipase (*LPL)* and Peroxisome Proliferator Activated Receptor Gamma *(PPARγ)*. Hypoxanthine Phosphoribosyltransferase 1 *(HPRT1)* and Tyrosine 3-Monooxygenase/Tryptophan 5-Monooxygenase Activation Protein Zeta (*YWHAZ*) were used as reference genes as recommended in [26]. All primers were available from an *in house* primer library (See Supplementary Table 1).

Gene expression was measured on diluted cDNA samples, in a Mx3005P qPCR System (Agilent Technologies) using QuantiFast SYBRGreen Master Mix (Qiagen) following the manufacturers recommendations. qPCR results were preprocessed using GenEx Pro. v7.0 (MultiD). Briefly, qPCR data was corrected by PCR efficiency, normalized using reference genes, relative quantities related to lean T0 group mean were calculated and data was log2 transformed. One-way ANOVA (Tukey-Kramer’s pairwise comparisons, 95% CI, *P*<0.05) was applied for statistical analysis in order to compare cell cultures across time (T0 and T21) and obesity-status (lean and obese).

## Results and Discussion

### Morphology, cell size and growth kinetics

After initial thawing, both lean and obese cell cultures exhibited very heterogeneous SVF populations. After 3-4 passages, most morphologies were lost and cultures presented largely homogenous morphologies. Lean samples showed a majority of cells with a long, straight morphology (Figure 1A); while obese samples showed a tendency towards a round, jagged morphology (Figure 1B), persisting throughout the passaging and differentiation processes. The surviving cell types were possibly selected due to: e.g. superior growth rate, better adaptation to the changed environment and nutrient source, better resistance or recovery from cryopreservation, trypsination time and deselection of cells lacking plastic-adherence. Despite several morphologies prevailing, the predominance of specific cell types in lean and obese cell cultures respectively indicates an innate difference between ASCs depending on the obesity status of the donor (lean vs obese). This could stem from a majority of cells of specific characteristics having been present in the SAT of the donor, or of different specific cell types from SAT of lean and obese donors, adapting better to the changed environment. Contrastingly, a study using SAT from obese and lean mice exhibited a similar mesenchymal phenotype for both lean and obese ASCs [32].

**Figure 1.**
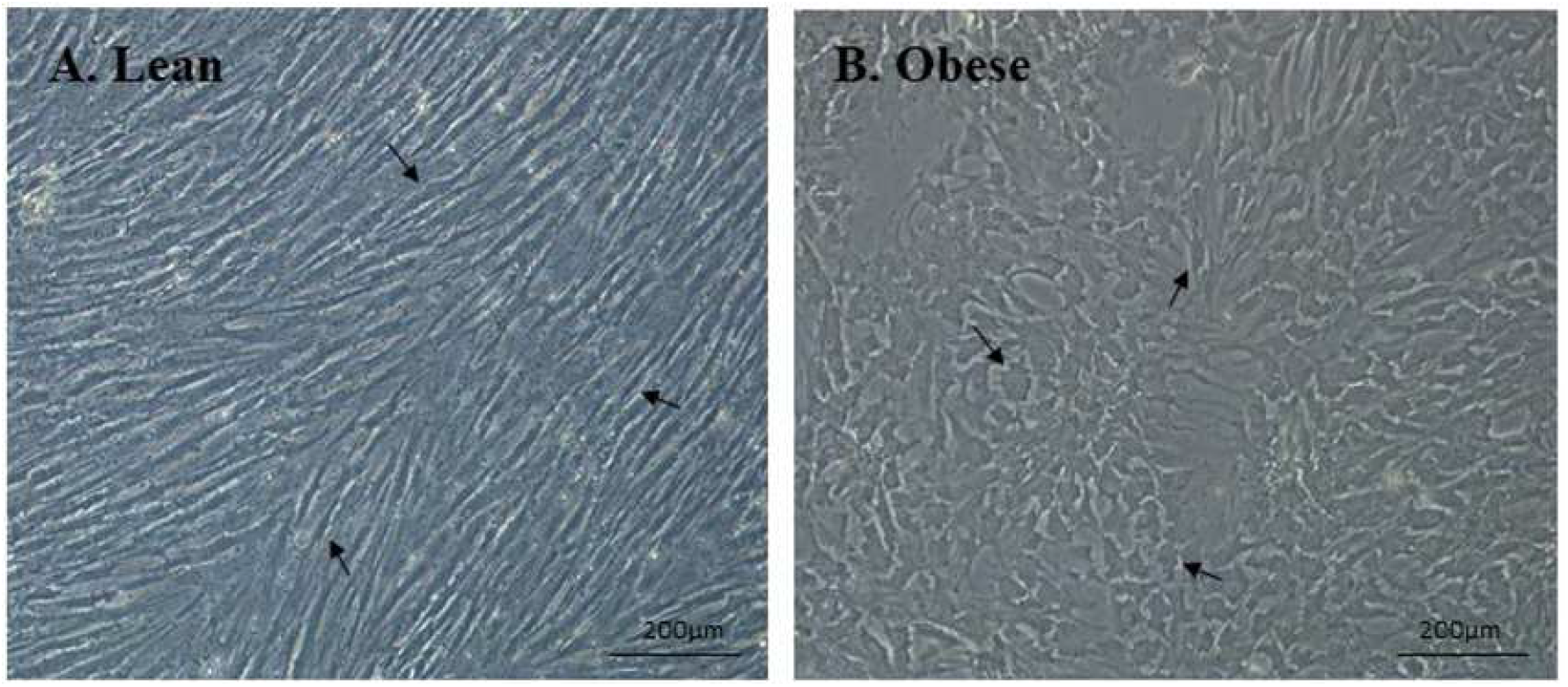
Lean and obese cells. After 1-2 passages, the majority of cells displayed a morphology characterized by a long, straight morphology (lean cultures) or a round, jagged morphology (obese cultures). Examples of lean and obese cells respectively is marked by arrows. 10x magnification, bars indicate 200μm.

During initial ASCs culturing, obese cell cultures showed more cells compared to lean cultures. This difference in cell number tended to disappear at later passages and lean and obese cultures exhibited largely similar viability, size and growth rate after first passage (Supplementary table 2) in agreement with another pig study [18]. In human and mouse studies, results are contradictory: the proliferation capacity of obese ASC cultures has been seen either increased [33, 34] or decreased compared with that of ASCs isolated from lean donors [10, 12, 35–37].

### Differentiation capacity

We differentiated two of the obese cultures (1 culture, F2, was excluded due to contamination during culturing) and the 3 lean cultures into the trilineages using specific media optimized for pig cells (see Supplementary text). Both lean (designated C1, C2 and C3) and obese cultures (designated F1 and, F3) exhibited capacity for trilineage differentiation (Supplementary Figure S1) and plastic adherence (results not shown), as described for mesenchymal stem cells [4]. We did not perform assessment of surface protein expression, to check compliance with MSC expression patterns, and therefore cannot state our cultures to be MSCs, despite MSC-like abilities, which should be taken into account.

The focus of the present study was the adipogenic differentiation. Therefore, we performed an optimization step of the adipogenic differentiation media for pig cells (data not shown). Specifically, we tried two different strategies according to Kim *et al*. (2016) and Zimmerlin *et al*. (2010) [29, 30] (see Supplementary text). After 21 days of adipogenic induction, the media from Zimmerlin *et al*. [29] showed more cells with lipid accumulation, in addition to higher cell viability and was subsequently used for our adipogenic differentiation procedure.

Both lean and obese cells started developing lipid droplets 5-7 days after induction, showing no difference in time of response. Nevertheless, obese cultures developed lipid-accumulating cells much more rapidly and, by the end of the differentiation period, greatly outnumbered lean cultures (see Figure 2A for a scoring of the adipogenic capacity). This result is in agreement with another pig study where they observed that ASC trans-differentiation towards adipocyte and osteocyte-lineages in obese ASCs was enhanced compared with lean ASCs [18]. In our study one lean culture (C3) was likewise advancing rapidly, but the remaining two lean cultures (C1 and C2) showed limited progression beyond day 7. After 21 days of adipogenic differentiation both obese cultures (F1 and F3) exhibited advanced adipogenesis, with extensive lipid accumulation (Figure 2A). Contrastingly, lean cultures diverged in their adipogenic capacity with one culture showing moderate adipogenesis, while the remaining two lean cultures showed very limited lipid accumulation. Optical measurement of the eluted Oil Red O confirmed the visual inspection, with all differentiated cultures showing higher absorbance than control wells, and absorbance of obese cultures exceeding that of lean cell cultures (Figure 2B).

**Figure 2.**
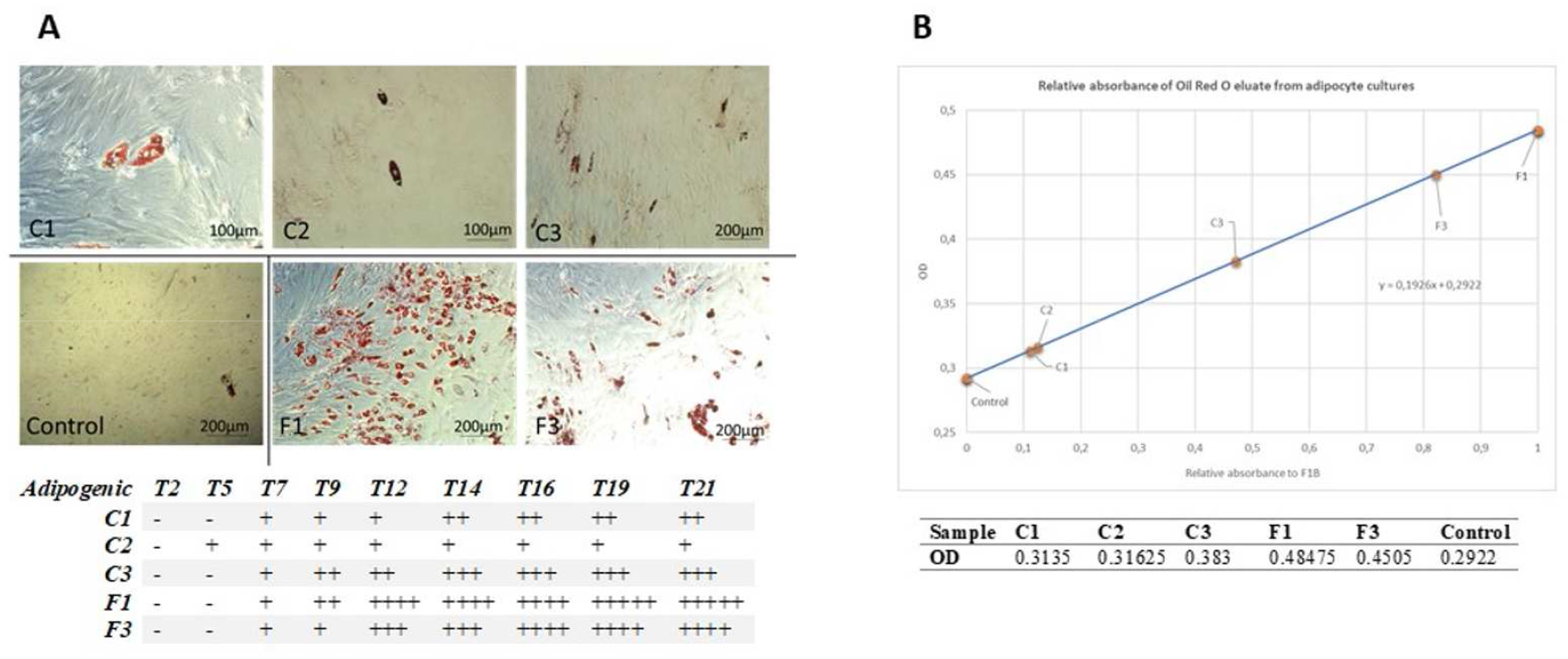
Adipogenic capacity of lean and obese cultures. A: Lipid droplets stained by Oil Red O. Representative images from Lean (C1, C2, C3), Obese (F1, F3) and Control cultures during adipogenic differentiation, following Oil Red O staining. C3, F1, F3 & Control: 10x magnification, bars indicate 200μm. C1 & C2: 20x magnification, bars indicate 100μm. Scoring of adipogenic capacity is given in reference to other cultures and timepoints. A score of 1+ is given upon first lipid formation, while 5+ signifies widespread lipid formation throughout the wells. T(n) indicate day n after adipogenic differentiation is initiated. B: Relative absorbance of eluted Oil Red O in isopropanol at 490nm. Culture values are averaged from adipogenic wells (n=4). Control value is averaged from all control wells (n=5).

These results seem to point towards a pre-disposition for adipogenic development in cultures originating from obese pig donors, not observed in the lean cultures. It is important to note, that the obese cultures originated from obese female GM and the lean cultures originated from lean males GM, and therefore a gender effect on differentiation cannot be dismissed. Studies on sex differences in adipogenesis are very limited, nevertheless, we consider the obesity-state as the most likely explanation for the observed differences. Human and rodent studies using ASCs isolated from white adipose tissue of obese individuals show conflicting results, some showed lower differentiation capacity to adipocytes in obese samples than their lean counterparts while other studies show no significant effect of the obesity status on differentiation capacity (for a review see [38]). Therefore, more research into this matter is warranted.

### Gene expression in cell cultures

We investigated gene expression for four adipogenesis-related genes (*LPL, ADIPOQ, PPARG, KLB*) in cell cultures derived from obese and lean SAT before (T0) and after 21 days of adipogenic induction (T21). Results showed a high divergence of gene expression between lean samples while obese samples were more similar, in agreement with the patterns observed on differentiation capacity (Figure 3).

**Figure 3.**
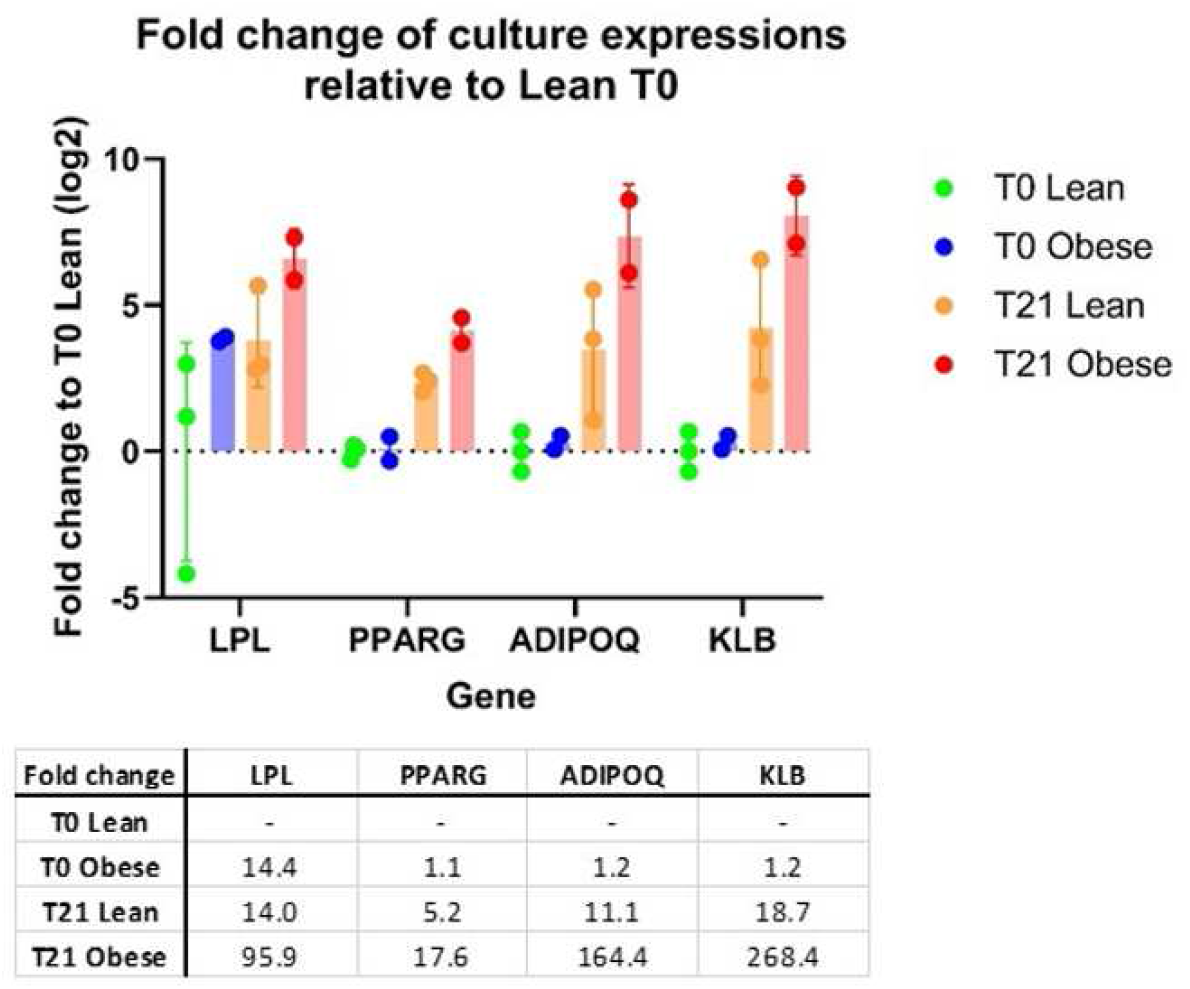
Relative expression (in log2) of adipogenic genes in cell cultures; all measurements are relative to Lean T0. Average fold changes for each gene in relation to Lean T0 cultures are shown.

In T0 cell cultures *LPL* showed tendency towards up-regulation in obese cultures relative to lean cultures (Fig. 3). Because LPL is involved in adipocyte metabolism, this result could suggest a level of adipogenic differentiation being initiated prior to induction and an inherent difference between lean and obese ASCs, also observed in other studies in human, mouse and pig [10–15, 18, 39, 40]. This result adds to our hypothesis, that obese cultures may be predisposed for adipocyte differentiation, causing more determined adipocyte development. *PPARG, ADIPOQ* and *KLB* were not expressed at measurable values in T0 cultures. Nevertheless, upon adipogenic induction, expression of all four genes increased in both lean and obese cultures. Indeed, after 21 days of adipogenic differentiation, the four genes showed tendency towards up-regulation (only *PPARG* was significantly up-regulated) in obese cultures relative to lean cultures with relevant fold changes (see Fig. 3). These four genes are commonly used adipogenic markers of ASC differentiation [39, 41], confirming the differentiation of both lean and obese cell cultures. *PPARG* is a master regulator of adipocyte differentiation controlling lipid storage and has been implicated in the pathology of numerous diseases including obesity [42]. ASCs from high fat diet-fed rats show reduced peroxisome proliferator-activated receptor-g (*PPARG*) expression which could affect ASC function [43]. *LPL* is involved in lipid metabolism and transport by catalyzing the rate-limiting hydrolysis of the triglyceride (TG) [44]. Adiponectin is an adipokine involved in the control of fat metabolism and insulin sensitivity [45]. KLB regulates indirectly bile acid synthesis by repressing cholesterol 7-alpha-hydroxylase (*CYP7A1*). *KLB* up-regulation in murine muscle-derived adipogenic progenitors has previously been linked to adipogenic differentiation [46]. This transcript showed the higher fold changes, even though it was not statistically significant due to the low sample number, supporting the notion of a more advanced adipogenic expression profile of obese ASCs upon differentiation (Fig. 3).

In summary, our results indicate inherent differences between lean and obese cell cultures that survive multiple cell passages, cryopreservation and differentiation *in vitro*. This points towards a cellular memory of the obesity status of the donor, priming ASCs derived from obese subjects towards an adipogenic fate. These inherent differences could be hypothesized to stem from e.g. epigenetic modifications. Indeed, methylation in promotors of genes with key role in adipogenesis (i.e. *PPARG*) have been associated with adipogenesis and differentiation, obesity, lipid metabolism, and adipose tissue expandability [47]. Relevant animal models are of key importance to study the properties and potential of stem cells derived from obese patients for possible future applications in human medicine. Further experiments investigating epigenetic marks in lean and obese ASC before and after differentiation could shed light into this hypothesis. Moreover, studies including several more biological replicates for each obesity status and taking gender into consideration would help to confirm the present findings.

## Conclusions

Despite the limited number and gender differences of the samples used in this study, our observations revealed intrinsic differences between lean and obese GM-derived ASC cells, with respect to morphology and differentiation capacity. In a context of cell therapy, this might convey, that stem cells from obese donors could display different therapeutic properties than those from lean donors, which cannot be neglected. However, the manner in which ASC function changes during obesity, and whether an inflammatory milieu in adipose tissue affects adipogenic differentiation, still remains controversial. Further investigations are warranted in order to progress in the ASC usage for cell-therapy in obese patients. In this context, we have shown that GM might constitute a valuable model in hand for further ASC research with special focus on investigating the effect of obesity in ASCs properties and functions.

## Acknowledgements

We would like to thank Henrik Duelund Pedersen from Ellegård Göttingen Minipigs and Trine Pagh Ludvigsen from Novo Nordisk for the donations of adipose tissue samples. We will also like to thank Tina B.N. Mahler and Minna B. Jakobsen for technical assistance.

## Author’s contribution

EØ, MF and SC conceived and designed the study. SC and MF accomplished the sample collection. MMM and SC did the laboratory work. MMM, EØ and SC contributed to the analysis and interpretation of data. MMM, SC and MF wrote the manuscript. All authors revised critically and approved the manuscript.

