## Supplementary material for "“Investigating the effect of obesity on adipose-derived stem cells (ASCs) using Göttingen Minipigs”"

**Table S1.** Growth rate, viability and size of cell cultures. Growth rate was determined as days from passage to confluent, upon passage from P(n) to P(n+1). Viability and live cell sizes were measured on an Automated Cell Counter upon passage from P(n-1) to P(n). P-values were calculated by student T-test (2-tail, homoscedastic).

| Growth rate (days) | Lean | Obese | P-value | Viability (%) | Lean | Obese | P-value | Live cell size (µm) | Lean | Obese | P-value |
|---|---|---|---|---|---|---|---|---|---|---|---|
| P1 | 8 | 3 | | P1 | 93.0 | 66.5 | | P1 | 16.97 | 11.05 | |
| | 7 | - | | | 86.5 | 80.5 | | | 18.79 | 13.01 | |
| | 4 | 4 | | | 84.0 | 90.5 | | | 20.20 | 15.26 | |
| Average | 6.33 | 3.5 | 0.174 | | 87.8 | 79.2 | 0.310 | | 18.65 | 13.10 | 0.022 |
| P2 | 7 | 5 | | P2 | 93.0 | 92.5 | | P2 | 16.05 | 19.84 | |
| | 10 | - | | | 91.5 | - | | | 17.04 | - | |
| | 4 | 7 | | | 87.5 | 93.0 | | | 23.01 | 19.26 | |
| Average | 7 | 6 | 0.700 | | 90.7 | 92.8 | 0.400 | | 18.70 | 19.55 | 0.782 |
| P3 | 5 | 7 | | P3 | 91.0 | 82.5 | | P3 | 17.51 | 15.93 | |
| | 7 | - | | | 88.0 | - | | | 19.10 | - | |
| | 5 | 5 | | | 92.5 | 91.0 | | | 19.37 | 17.37 | |
| Average | 5.67 | 6 | 0.789 | | 90.5 | 86.8 | 0.374 | | 18.66 | 16.65 | 0.117 |

**Table S2**. Primers used for gene expression profiling. Ampliqon size in bp.

| Gene name | Forward primer | Reverse primer | Ampliqon size |
|---|---|---|---|
| ***ADIPOQ*** | CGAGAAGGGTGAGAAAGGAG | TAGGCGCTTTCTCCAGGTT | 123 |
| ***LPL*** | CCCTGGCTTTGCTATTGAGA | ACTTGTCGTGGCATTTCACA | 139 |
| ***PPARG*** | GCCGTGTCTGTGGGGATAAA | CCGACAGTTAAGATCGCACCT | 128 |
| ***KLB*** | TGGTTCACAGACAGTCACGT | TGCCATTCAAAGCCATCCAG | 152 |
| ***HPRT1*** | GGACTTGAATCATGTTTGTG | CAGATGTTTCCAAACTCAAC | 91 |
| ***YWHAZ*** | TGATGATAAGAAAGGGATTGTGG | GTTCAGCAATGGCTTCATCA | 203 |

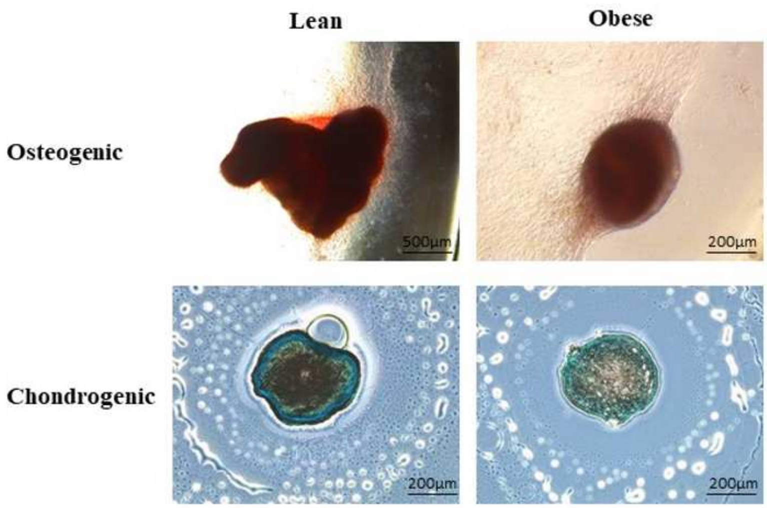


**Figure S1**. Representative images of osteogenic and chondrogenic differentiation in lean and obese cultures. Osteogenic cultures stained with Alizarin S Red. Osteogenic, lean: 4x magnification, bar indicate 500µm. Osteogenic, obese: 10x magnification, bar indicate 200µm. Chondrogenic cultures stained with Alcian Blue: 10x magnification, bars indicate 200µm.

**Supplementary text. Differentiation media:**

**Adipogenic media:** DMEM/F-12 (Sigma-Aldrich) +10% FBS (Gibco) + 1% Penicillin-Streptomycin (Thermo Fisher Scientific), 1µM dexamethasone (Sigma-Aldrich), 0.5µM 3-isobutyl-1-methylxanthine (IBMX)(Sigma-Aldrich), 60µM indomethacin (Sigma-Aldrich), 10µg/ml insulin solution from bovine pancreas (Sigma-Aldrich).

**Osteogenic media**: DMEM/F-12 (Sigma-Aldrich) + 10% FBS (Gibco) + 1% Penicillin-Streptomycin (Thermo Fisher Scientific), 10nM dexamethasone (Sigma-Aldrich), 10mM β-glycerophosphate (Sigma-Aldrich), 50µg/ml ascorbic acid (Sigma-Aldrich).

**Chondrogenic media**: DMEM/F-12 (Sigma-Aldrich) w. 1.35mg/ml glucose + 1% Penicillin-Streptomycin (Thermo Fisher Scientific), 100nM dexamethasone (Sigma-Aldrich), 10µl/ml insulin-transferrin-sodium selenite media supplement (ITS)(Sigma-Aldrich), 1mM sodium pyruvate (Gibco), 50µg/ml ascorbic acid (Sigma-Aldrich), 5ng/ml transforming growth factor β1 (TGF-β1) from porcine platelets (Sigma-Aldrich)(100x solution used = 1µg TGF-β1 dissolved in 1ml TGF-β solvent (4 nM HCl + 100µl bovine serum albumin (BSA)(Sigma-Aldrich) in 1ml PBS)).

Prior to adipogenic differentiation in porcine cells, we tested 2 different adipogenic media, as described by Kim *et al.* (2016) and Zimmerlin *et al.* (2010). After 21 days of adipogenic induction, the media from Zimmerlin *et al.* showed more cells with lipid accumulation, in addition to higher cell viability and was used for our differentiation procedure.

Two of the lean cell cultures were subjected to an additional passage before differentiation, due to insufficient cell numbers. No visible difference between P3 and P4 lean cultures could be determined in differentiation procedures.
